# Behavioural outcomes of adult female offspring following maternal stress and perinatal fluoxetine exposure

**DOI:** 10.1101/137414

**Authors:** Veronika Kiryanova, Sara J. Meunier, Richard H. Dyck

## Abstract

Depression, anxiety, and stress are common in pregnant women. One of the primary pharmacological treatments for anxiety and depression is the antidepressant fluoxetine (Flx). Maternal stress, depression, and Flx exposure are known to effect neurodevelopment of the offspring, however, their combined effects have been scarcely studied, especially in female offspring. The present study investigated the combined effects of maternal stress during pregnancy and perinatal exposure to Flx on the behaviour of female mice as adults. METHODS: Mouse dams were exposed to either chronic unpredictable stress (embryonic (E) day 7 to E18), or FLX (E15-postnatal day 12), or a combination of stress and FLX or left untreated. At two months of age, the female offspring went through a comprehensive behavioural test battery. RESULTS: Maternal stress led to increased activity and alterations of prepulse inhibition in the adult female offspring. Maternal treatment with Flx had a potentially beneficial effect on spatial memory. The combination of prenatal stress and perinatal Flx exposure did not interact in their effects. These results suggest that gestational Flx exposure may have a limited negative impact on female offspring.

## 1. Introduction

Stress-related disorders such as depression and anxiety are common in pregnant women: 10 - 18% of pregnant women present with symptoms of depression, and 6-14% present with symptoms of anxiety disorders [1-4]. This prevalence is even higher in pregnant women with lower socioeconomic status due to, in part, increased frequency of stressful life events [5-8]. Depression is a very stressful experience, it impacts a woman’s sleep, eating, and self-care and the level of care she can provide to her child. Stress, depression, and anxiety during pregnancy can also lead to adverse obstetric outcomes and affect fetal and neonatal development and behaviour [9, 10].

Primary pharmacological treatments for anxiety and depression are selective serotonin reuptake inhibitors (SSRIs) [11]. Out of the 6% of pregnant women who are prescribed an SSRI at some point during their pregnancy, between 20-25% will receive fluoxetine (Flx; brand names Prozac, Sarafem, Rapiflux) [12].

Maternal intake of antidepressants and maternal anxiety, depression, and stress can affect obstetric outcomes and child’s development and behaviour. Early exposure to these environmental events are linked with a decrease in gestational length and a reduction in birth weight [13-15], and may lead to changes in emotional behaviour, including anxiety, in children [16, 17]. Beyond the childhood years, effects of maternal anxiety, depression, and stress and maternal Flx exposure are not yet fully understood. Animal studies have been essential in uncovering these effects [for a review see 18, 19].

Animal studies utilize maternal stress as a model of maternal depression and anxiety. Studies show that maternal stress and maternal exposure to Flx both have long-term effects on the mouse and rat offspring. These maternal experiences alter offspring’s learning and memory (stress: [20-25]; Flx: [26-28]), aggression levels (stress: [29-32]; Flx: [28, 32-34]), circadian behaviours (stress: [35]; Flx:[36]), and depressive-and anxiety-like behaviours (stress: [25, 37-39]; Flx: [26-28, 40-43]).

While necessary, examining the offspring of healthy animals is of minor ethological validity when attempting to draw inferences about exposed humans. This is because Flx exposure in humans does not typically happen in isolation; Flx exposure is concurrent with maternal stress, anxiety, and/or depression. Animal studies are beginning to examine combined effects of perinatal stress and Flx exposure on adult offspring outcomes. Some studies demonstrate that Flx and stress have specific long-term effects on the offspring. They have distinct effects on reproductive behaviour [44, 45], sexual brain differentiation [44], and depressive-like behaviour [46] in rats, and anxiety-like behaviour [32, 46] and hippocampal and cortical BDNF signaling in rats and mice [32, 46]. Other studies demonstrate that while maternal exposure to stress and Flx can have distinct effects, treatment with Flx early in life can counteract some of the effects of maternal stress. In mice, these include alterations in spatial learning [47], HPA-axis reactivity [48], aggressive behaviours [32], and circadian behaviours [35], while in rats, it includes sensitivity to post-operative pain [49]. In both rodent models, Flx counteracts stress-induced alterations in hippocampal morphology [47, 50]. However, most behavioural studies were conducted in male offspring; behaviours of adult female offspring have not yet been examined aside from reproductive [45] and depressive-like behaviours of rats [46]. A thorough assessment of outcomes in female offspring is warranted. Past studies provided indications that outcomes of female offspring may differ from that of males. First, behavioural baselines show sexual dimorphism [51-53]. Second, male and female animals are differently affected by stress [54-57] and Flx administration [58-60]. And, most importantly, male and female mice show different patterns of behavioural changes following maternal stress [22, 23, 61-65] and perinatal Flx administration [43, 63, 66].

The goal of the present study was to investigate the effects of prenatal stress and effects of perinatal exposure to Flx, separately and in combination, on the behaviour of the adult female offspring. This study sought to provide a more complete insight by conducting an in-depth behavioural analysis of the offspring’s cognitive ability, memory, anxiety, sensorimotor information processing, and exploratory and risk assessment behaviours.

## 2. Methods

### 2.1 Animals

All mice were kept on a 12:12-h light/dark (LD, 1,500 lx/0 lx) and were provided *ad libitum* access to food (LabDiet Mouse Diet 9F, #5020) and water throughout the experiment with the exception of stress manipulation measures (see below). Animal treatment and husbandry were performed in accordance with the Canadian Council on Animal Care. All experimental procedures were approved by the Ethics Committee for Animal Research at the University of Calgary.

C57BL/6 breeders were obtained from the University of Calgary Biological Sciences breeding facility (Calgary, AB, Canada) and the Charles River animal facility (Wilmington, MA, USA; equally dispersed between treatment groups). At 3 months of age, male and female mice were paired in breeding triads (one male and two females) for a duration of 4 days. The day the seminal (copulation) plug was first observed was considered embryonic (E) day zero. The day of birth was considered to be postnatal (P) day 0.

Pregnant dams were randomly assigned into four groups: (I) dams untreated (CON; n=6); (II) dams subjected to chronic unpredictable stress (PS; n=6); (III) dams administered Flx (FLX; n=5); (IV) dams subjected to PS and administered Flx (PS+FLX; n=6). One day following parturition litters were culled to 8 pups (pups that were culled were picked randomly). Litters were housed together with their mother until weaning at P21 after which they were housed with same sex littermates in groups of 3-5. Final groups for behavioural testing consisted of 9 female mice per group; no more than 2 female pups were used from each litter. Male offspring were used in a different set of experiments.

### 2.2 Chronic Unpredictable Stress Paradigm

From E4 to E18, pregnant dams were subjected to a regimen of chronic unpredictable stressors (PS). The PS paradigm was modified from that used by Grippo and colleagues [67] and used as described in [32] and [35]. Stressors included restricted access to food (5- and 6-hour food deprivation; delivered during the day and overnight), continuous lighting overnight, cage tilt (home cage tilted 30⁏; delivered overnight or for 7 hours during the day), paired housing (animals were paired with another pregnant dam in either their home-cage or the other dam’s cage, assigned in a counterbalanced fashion; delivered overnight), foreign object in cage (a novel plastic or glass object; overnight), soiled cage (100ml of clean water spilled on bedding; overnight), irregular 16kHz tones (played at 80dB; delivered for one hour during the day), white noise (played at 80dB; delivered for 3 hours during the day), and restraint (for the duration of two hours; delivered three times during the stress regimen). At least one and at most four stressors were delivered each day.

### 2.3 Drug Treatment

Between E15 and P12, pregnant dams were administered fluoxetine hydrochloride (Flx; Sigma-Aldrich, Saint Louis, MO, USA) in their drinking water at a dose of 25mg/kg/day as previously described [28]. The Flx concentration in the water was based on the mouse’s weight at the time and the mouse’s water consumption for the preceding 48 hours. This concentration was re-calculated every 48h to control for weight gain during the gestation period. This treatment protocol has been demonstrated to lead to pup brain levels of Flx and its metabolite norfluoxetine that fall within the range observed in post-mortem brain tissue of humans who took this medication [28].

### 2.4 Measurement of Fluoxetine and Norfluoxetine in Pup Brains

#### 2.4.1 Sample collection

Pup brains were collected at P12 (FLX: n=4, from 3 litters; PS+FLX: n=5, from 4 litters). After decapitation the brain was removed, the spinal cord was severed posterior to the cerebellum, and the olfactory bulbs were removed. Cortical structures (cortex and hippocampus) were peeled away from the thalamus and the subcortical portion was bisected between the cerebellum and inferior colliculus. The three resulting brain areas were separately frozen on dry ice and analyzed as: i. cortex and hippocampus, ii. thalamus/midbrain, and iii. hindbrain/cerebellum.

#### 2.4.2 Sample analysis

The sample preparation method was based on Raap et al. [68]. The concentrations of fluoxetine and norfluoxetine were quantified by high-performance liquid chromatography. Sample preparation and analysis were performed exactly as previously described [28].

### 2.5 Behavioural Analysis

A battery of well-validated behavioural tests was used to assess various patterns of adult behaviour in mice. The behavioural test battery we used was based upon previous reports assessing long term behavioural consequences of both stress and Flx treatments. Starting at 2 months of age, the offspring were administered behavioural tests assessing exploratory behaviour and locomotion, anxiety-related behaviours, sensorimotor gating and acoustic startle response, and spatial and fear memory.

Tests were performed in the following order: Open field and the elevated plus maze administered to all groups. The next three tests (Morris water test, passive avoidance, and prepulse inhibition) were conducted in a counterbalanced fashion, in that equal number of animals from each group received these tests in a different order. All testing was conducted during the light phase of the light-dark cycle, between 9am and 5pm. Tests were conducted five to seven days apart.

#### 2.5.1 Open Field

The open field task assesses an animal’s movement in a large brightly lit, walled arena. By analyzing speed and distance travelled, differences in locomotor activity between groups can be measured and a comparison of time spent adjacent to the arena walls (thigmotaxis) and time spent in the arena’s center provide a relative measure of anxiety [69].

In the current experiment a 1.2m diameter brightly lit (170±30lux) white Plexiglas arena surrounded by a 35cm high wall was used. Mice were placed individually in the arena’s center and allowed to explore for 5 min. Locomotor activity was recorded using an overhead mounted camera and HVS Image 2020 Plus software (HVS Image Ltd, Twickenham, Middlesex, UK). Thigmotaxis (Percent of time spent in the outer third of the arena), total distance traveled, and speed were measured. Following each trial the arena was cleaned using a 70% ethanol solution to remove olfactory cues.

#### 2.5.2 Elevated Plus Maze

The elevated plus maze test was conducted as previously described [28, 70, 71]. The elevated plus maze (EPM) test is a well-validated measure of anxiety-like behaviour in rodents [72]. The plus-shaped platform consists of two opposing enclosed arms (5cm × 28cm) and two opposing open arms (5cm × 28cm) extending outward, at right angles, from a central platform (5cm × 5cm) elevated 28cm above the floor; the platform was illuminated evenly by an overhead light (75±8 lux). Animals were placed individually in the maze center facing an open arm then allowed to freely explore for 5 min.

The percent of total time a mouse spent on the open arms, in the center or on closed arms, as well as the number of entries into open and closed arms, speed and distance travelled were measured using ANY-maze software version 4.73 (Stoelting Company, Wood Dale, Illinois, USA). Arm entry was operationally defined as 4 paws within an arm, or 95% of animal within an arm. Scanning behaviour (head dip), instances on the open arms when the animal looked over the edge towards the floor, and risk assessment behaviours (protective stretch), instances in which the animal stretched its head and forepaws into an open arm from a closed arm or the center space without making a full entry into the open arm were scored manually by the researcher [73]. The number of entries into open and closed arms, speed, distance, and time spent active were also taken as a measure of locomotor activity [74].

#### 2.5.3 Morris Water Task

The Morris water task was conducted as previously described [32]. Briefly, MWT pool was white in color, 1.2 m in diameter and filled with water (19.5 ± 0.5° C) to a depth of 20 cm. A clear plexiglas, square platform (20 × 16) was positioned in the center of one quadrant with the surface 1.5cm below the water’s surface. The water was made opaque by adding skim milk powder. Animals were video recorded from overhead using HVS Image 2020 Plus software (HVS Image Ltd, Twickenham, Middlesex, UK).

During the first four training days, the mice were subjected to four trials each day, where they were placed into the pool facing the wall at one of the cardinal compass points, chosen in a pseudo-random order (N/S/E/W). Each trial lasted 60s and was followed by a 15s learning period where the mouse stayed atop the platform. Trials were separated by a 7-minute inter-trial interval. Trials were analyzed for speed, distance travelled, and latency to find the platform.

On the fifth day (probe trial) the platform was removed from the pool and animals were placed in a novel start location and given 60s to swim freely. The initial latency to where the platform had been training days was recorded. Furthermore, the number of times the mouse swam through the area of platform’s previous location (across or within one body space) was counted and analyzed.

#### 2.5.4 Passive Avoidance

Animals were placed individually in a white Plexiglas chamber with a metal grate floor (20.32cm × 25.4cm × 35.71cm), part of the dual-chamber LM1000-B avoidance system (Hamilton-Kinder LLC, San Diego, CA). Following acclimatization (2 min) the chamber door opened, allowing animals to enter a second dark chamber (black walls and roof) with the same metal grate floor. When the mouse had all four paws within the dark chamber, it was administered a mild foot shock (0.5mA, duration 1s) through the metal floor. Twenty-four hours later the animals were placed individually within the white chamber with the door to the dark chamber already opened. Latency to enter the dark chamber was recorded for a maximum of 3 min.

#### 2.5.5 Prepulse Inhibition

Prepulse inhibition (PPI) is a measure of the magnitude of acoustic startle response to a stimulus when it is preceded by a weaker, sub-threshold stimulus (prepulse). PPI is a well validated measure of attentional impairments, sensory processing, and sensorimotor gating, known to be disrupted in various forms of psychopathology (Crawley, 2007). PPI and acoustic startle response were analyzed using the SM100SP Startle Monitor system (Hamilton-Kinder LLC; San Diego, CA, USA). The apparatus consisted of a sound-attenuated chamber (27.6cm × 35.6cm × 49.5cm) housing a clear plexiglas chamber (10cm × 3.8cm) with a piezoelectric motion monitor. Mice were placed into the chamber, provided a 5 min acclimatization period with a 65 dB steady ambient noise presented. Following this acclimatization period, a habituation phase took place in which a sound of 120dB (duration 40ms) was delivered 10 times every 5-20s. Following habituation, the testing phase occurred where 3 types of trials were each presented 10 times separated by 5-20s intervals. The trials consisted of (i) no pulse, (ii) pulse alone (120dB, duration 40ms), and (iii) prepulse (80dB, duration 20ms) followed by a pulse (120dB, duration 40ms; 100ms interval). Percent PPI was calculated using the formula: 100-(100 × startle amplitude on prepulse trial)/startle amplitude on pulse alone trial. The animal’s average startle response to a pulse, as well as percent PPI, were analyzed.

#### 2.5.6 Fear Conditioning

Fear conditioning paradigms based on Pavlovian conditioning were used to assess learning and memory of a negative stimulus (foot shock). Mice experienced the pairing of a novel stimulus (tone) and context (chamber) with an adverse event (foot shock). Contextual and cued fear conditioned responses were assessed using a dual-chamber LM1000-B avoidance system (Hamilton-Kinder LLC, San Diego, CA) following a testing procedure modified from that used by Saxe and colleagues [75], as detailed below. Experiments took place across 3 days and consisted of a conditioning day, a cued fear memory testing day, and a contextual fear memory testing day.

On the conditioning day, mice were placed within the conditioning chamber (20.32cm × 25.4cm × 35.71cm) for a 2-min habituation period. They then received 3 pairings, at 2min intervals, of 80dB tones (duration 20s; conditioned stimulus) that terminated with a mild foot shock (0.5mA, duration 1s; unconditioned stimulus). The following day, contextual fear conditioning was assessed. On this day the mice were placed back into the chamber in which they had experienced the initial pairings of shock and tone for 4min, and freezing behaviour (absence of motor movement aside from breathing) in response to the original conditioning context was assessed across the entire session. The mice experienced neither tone nor foot shock on the contextual fear conditioning day. On the third day mice were placed in an environmentally altered chamber (color of walls (black to white), scent of environment (neutral to coconut), disinfectant used (70% ethanol to Virkon), experimenter glove material (latex to nitrile), floor of chamber (from metal to plastic) to reduce contextual cues. Following a 2min habituation period, mice were exposed to an 80dB tone (duration 40s) that was not followed by a foot shock. Freezing behaviour in response to the conditioned stimulus (tone) was assessed one minute before the tone through the 40s duration of the tone. On the second and third days (contextual and cued fear conditioning) mice were recorded using an overhead camera (Sony DCR-TRV10), which were analyzed for freezing behaviour using ANY-maze software version 4.73 (Stoelting Company, Wood Dale, Illinois, USA).

### 2.6 Data Analysis

Tests that involved a single measurement (i.e., open field, elevated plus maze, prepulse inhibition) were analysed using a two-way (fluoxetine × stress) factorial analysis of variance (ANOVA). For tasks that involved repeated testing of the animals (e.g., passive avoidance, Morris water test, and levels of serotonin, Flx and NorFlx in different brain regions) a split plot ANOVA was computed. Tukey’s post hoc test for multiple comparisons was used to assess differences between individual treatment groups. Protected t-tests using the Bonferroni correction were used for multiple comparisons when following up main effects in repeated measurements (e.g., Morris water test). To examine whether litter effects contributed to any of the significant findings, all significant treatment effects were followed up (*p* < 0.05) with a nested ANOVA, this assessed the effect of litter nested within group. We failed to find any significant litter effects (*p* >.10) (data not shown). All statistics were two-tailed. Values of *p* < 0.05 were considered significant. All data and all figures are reported as means ± standard error of the mean (SEM).

## 3. Results

### 3.1 Actual Dose of Fluoxetine

Increased water consumption throughout pregnancy and lactation resulted in an actual mean Flx dose of 27.42±.55 mg/kg/day, instead of intended 25 mg/kg/day. Administered Flx dosage did not significantly differ between FLX and FLX+PS groups, and there was no significant group by day interaction.

### 3.2 Fluoxetine and Norfluoxetine in Pup Brains

In the P12 offspring that were exposed to fluoxetine, the mean brain fluoxetine level was 0.361±0.06 µg/g and the overall brain level of NorFlx was 3.73±0.33 µg/g. There was no significant difference among brain regions in fluoxetine or NorFlx concentration.

### 3.3 Behavioural Evaluation of Adult Animals

#### 3.3.1 Open Field

There was no effect of PS, Flx treatment, or PS by Flx treatment interaction on thigmotactic behaviour, speed and distance travelled in the open field (see Table 1A).

**Table 1A:** Summary of results of behavioural testing battery demonstrating effects of prenatal stress and perinatal fluoxetine exposure on behaviour of female mice as adults

| Behavioural Test | Control | Fluoxetine | Stress | Stress+<br>Fluoxetine | Main Effect of Flx | Main Effect of Stress | Flx and Stress Interaction |
| --- | --- | --- | --- | --- | --- | --- | --- |
| *Open Field* |  |  |  |  |  |  |  |
| Thigmotaxis time (%) | 60.35±4.02 | 69.48±4.27 | 53.03±6.60 | 56.78±5.75 | $F_{(1, 32)} = 1.49, p = .231$ | $F_{(1, 32)} = 3.61, p = .066$ | $F_{(1, 32)} = .26, p = .613$ |
| Speed (m/s) | .11±.001 | .11±.010 | .086±.002 | .11±.011 | $F_{(1, 32)} = 1.42, p = .242$ | $F_{(1, 32)} = 3.20, p = .083$ | $F_{(1, 32)} = 1.15, p = .291$ |
| Distance (m) | 33.29±3.42 | 34.41±2.93 | 26.02±.96 | 32.12±2.93 | $F_{(1, 32)} = 1.75, p = .196$ | $F_{(1, 32)} = 3.06, p = .090$ | $F_{(1, 32)} = .828, p = .370$ |
| *Elevated Plus Maze* |  |  |  |  |  |  |  |
| Open arm time (%) | 10.87±3.12 | 8.71±1.34 | 14.29±4.51 | 14.09±3.77 | $F_{(1, 32)} = .12, p = .730$ | $F_{(1, 32)} = 1.67, p = .205$ | $F_{(1, 32)} = .08, p = .776$ |
| Open arm entry | 5.33±1.37 | 4.22±0.52 | 5.89±1.02 | 5.22±1.24 | $F_{(1, 32)} = .67, p = .421$ | $F_{(1, 32)} = .51, p = .481$ | $F_{(1, 32)} = .04, p = .840$ |
| Closed arm entry | 9.00±.86 | 11.22±1.63 | 11.67±1.04 | 12.56±1.41 | $F_{(1, 32)} = 1.49, p = .231$ | $F_{(1, 32)} = 2.46, p = .126$ | $F_{(1, 32)} = .27, p = .605$ |
| Speed (m/s) | .018±.001 | .021±.002 | .023±.001 | .023±.002 | $F_{(1, 32)} = .12, p = .731$ | $F_{(1, 32)} = 3.22, p = .082$ | $F_{(1, 32)} = .75, p = .392$ |
| Distance (m) | 5.57±.45 | 6.23±.70 | 7.07±.48 | 6.81±.63 | $F_{(1, 32)} = .12, p = .728$ | $F_{(1, 32)} = 3.27, p = .080$ | $F_{(1, 32)} = .63, p = .435$ |
| Time active (%) | 55.23±13.66 | 57.41±10.43 | 63.84±6.19 | 61.09±13.30 | $F_{(1, 32)} = .01, p = .941$ | $F_{(1, 32)} = 2.65, p = .113$ | $F_{(1, 32)} = .43, p = .517$ |
| Protected stretches | 10.67±1.37 | 11.22±0.86 | 9.44±0.77 | 11.22±1.23 | $F_{(1, 32)} = 1.15, p = .292$ | $F_{(1, 32)} = .315, p = .578$ | $F_{(1, 32)} = .315, p = .578$ |
| Head dips | 9.22±2.62 | 9.00±1.63 | 11.67±3.57 | 11.44±2.76 | $F_{(1, 32)} = .01, p = .936$ | $F_{(1, 32)} = .80, p = .379$ | $F_{(1, 32)} = .000, p > .999$ |
| *Prepulse Inhibition* |  |  |  |  |  |  |  |
| PPI (%) | 34.36±4.39 | 25.31±4.88 | 44.86±3.50 | 43.41±2.72 | $F_{(1, 32)} = 1.75, p = .195$ | $F_{(1, 32)} = 13.00, p = .001$ | $F_{(1, 32)} = .918, p = .345$ |
| Startle response | .39±.057 | .39±.058 | .40±.035 | .54±.044 | $F_{(1, 32)} = 1.97, p = .170$ | $F_{(1, 32)} = 2.88, p = .099$ | $F_{(1, 32)} = 1.59, p = .217$ |

*Note:* Significant results ( $p < 0.05$ ) are reported in bold font; n=9 per group. All data are
presented as mean ± SEM.

#### 3.3.2 Elevated Plus Maze

There was no effect of PS, Flx administration, or PS by Flx interaction on any measures on the EPM (see Table 1A).

#### 3.3.4 Passive Avoidance

It took longer for mice to enter the dark chamber on day two, compared to day one (*F*_(1, 32)_ = 119.10, *p* < .001) indicating that all mice learned the dark chamber/shock association (see Table 1B). The proportion of mice that entered the dark chamber on day two was not affected by perinatal Flx administration (𝒳^*2*^ (1, *N* = 36) = .423, *p* = .691), or by PS (𝒳^*2*^ (1, *N* = 36) = .423, *p* = .691).

**Table 1B:** Summary of results of behavioural testing battery demonstrating effects of prenatal stress and perinatal fluoxetine exposure on behaviour of female mice as adults

| Behavioural Test | Main Effect of Flx | Main Effect of Stress | Flx and Stress Interaction | Effect of Time |
| --- | --- | --- | --- | --- |
| *Passive avoidance* |  |  |  |  |
| Latency | $F_{(1, 32)} = .91, p = .347$ | $F_{(1, 32)} = 1.20, p = .282$ | $F_{(1, 32)} = 3.24, p = .081$ | $F_{(1, 32)} = 119.10, p < .001$ |
| *Morris Water Task* |  |  |  |  |
| *Training Days* |  |  |  |  |
| Latency | $F_{(1, 32)} = 6.13, p = .019$ | $F_{(1, 32)} = 1.01, p = .323$ | $F_{(1, 32)} = .002, p = .967$ | $F_{(3, 96)} = 81.37, p < .001$ |
| Speed | $F_{(1, 32)} = .03, p = .865$ | $F_{(1, 32)} = 5.04, p = .032$ | $F_{(1, 32)} = .34, p = .562$ | |
| Thigmotaxis | $F_{(1, 32)} = 1.63, p = .211$ | $F_{(1, 32)} = 1.46, p = .235$ | $F_{(1, 32)} = .06, p = .804$ | |
| *Probe Trial* |  |  |  |  |
| Latency | $F_{(1, 32)} = 4.18, p = .049$ | $F_{(1, 32)} = 0.96, p = .333$ | $F_{(1, 32)} = .33, p = .572$ | |
| Time Crossed | $F_{(1, 32)} = 9.98, p = .003$ | $F_{(1, 32)} = .17, p = .683$ | $F_{(1, 32)} = .17, p = .683$ | |

*Note:* Significant results ( $p < 0.05$ ) are reported in bold font; n=9 per group. All data are presented as mean ± SEM.

#### 3.3.5 Prepulse Inhibition

Female offspring of prenatally stressed dams had a higher PPI (44.1±9.2%) compared to offspring of non-stressed dams (29.8±14.3%; *F*_(1, 32)_ = 13.00, *p* = .001; Fig 1). There was no effect of perinatal Flx exposure or PS by Flx interaction on PPI. Furthermore, no effect of PS, Flx administration, or PS by Flx interaction was observed on the startle response.

**Figure 1.**
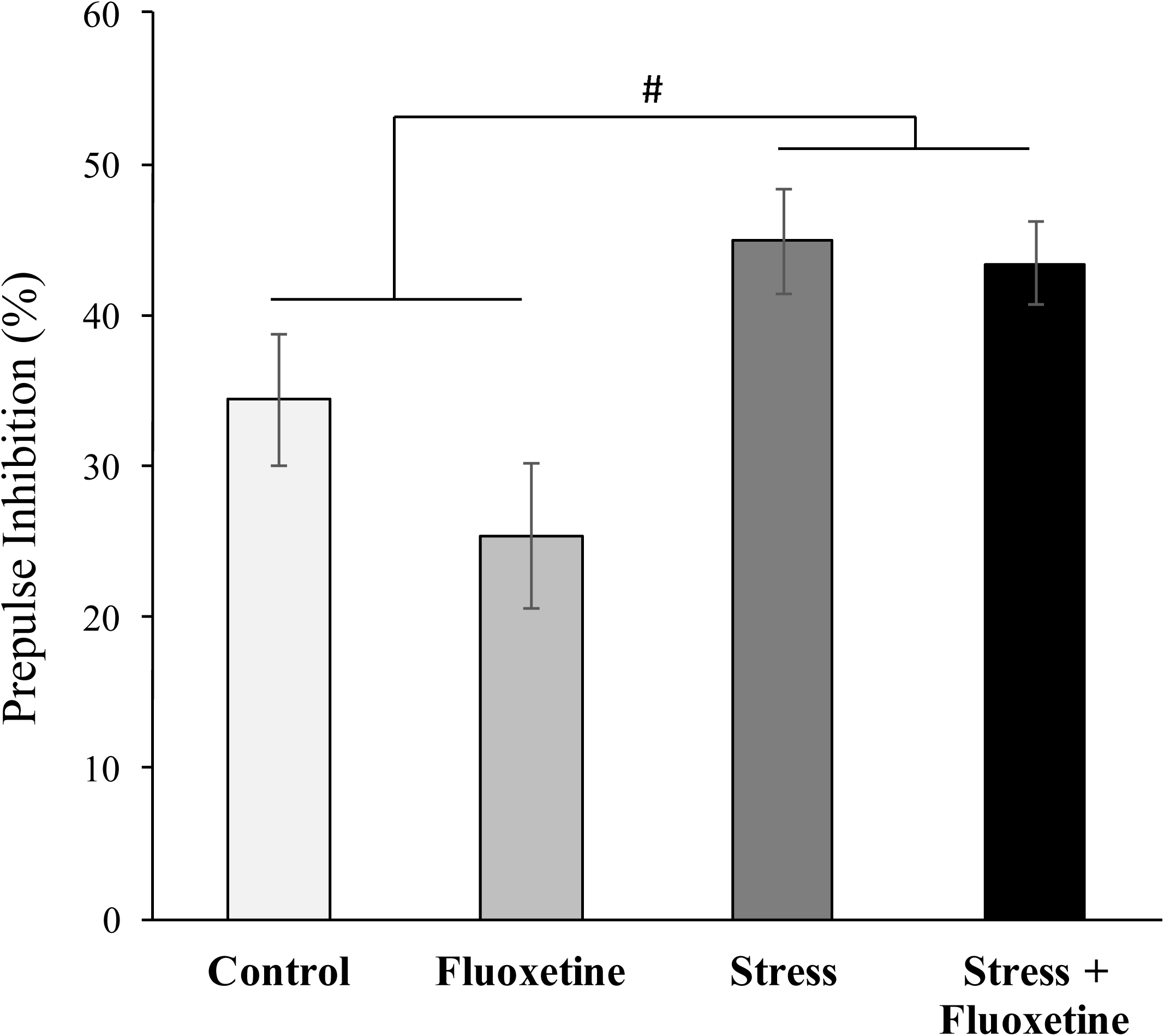
Prepulse inhibition. Adult female offspring of dams subjected to chronic variable stress show enhanced prepulse inhibition (n=9 per group); # denotes significant main effect of stress, *p* < 0.05. All data are presented as mean ± SEM.

#### 3.3.6 Morris Water Task

Latency to reach the platform decreased with training, indicating that all mice were able to learn the task (*F*_(3, 96)_ = 81.37, *p* < .001; Fig 2A). While there was a significant effect of Flx on latency to reach the platform during the training days (*F*_(1, 32)_ = 6.13, *p* = .019; Fig 2A), with Flx exposed animals reaching the platform slower than non-Flx exposed animals, after correcting for multiple comparisons, there was no significant difference between Flx exposed and non-exposed animals on latency to reach the platform on any of the training days (day1: *t*_(34)_ = 2.07, *p* = .046; day2: *t*_(34)_ = 1.58, *p* = .123; day3: *t*_(34)_ = 1.86, *p* = .071; day4: *t*_(34)_ = .75, *p* = .460). Female offspring of stressed dams swam faster (.176±.005m/s) than offspring of dams not subjected to PS (.161±.005m/s; *F*_(1, 32)_ = 5.04, *p* = .032; Fig 2B). During the probe trial, mice perinatally exposed to Flx crossed the platform area significantly more times (Flx-exposed:2.7±1.4; not exposed to Flx:1.5±.92; *F*_(1, 32)_ = 9.98, *p* = .003; Fig 2C) and found the platform faster (Flx-exposed:17.7±16.7s; not exposed to Flx:31.5±24.1s; *F*_(1, 32)_ = 4.18, *p* = .049; Fig 2D) than mice not exposed to Flx (see Table 1B).

**Figure 2.**
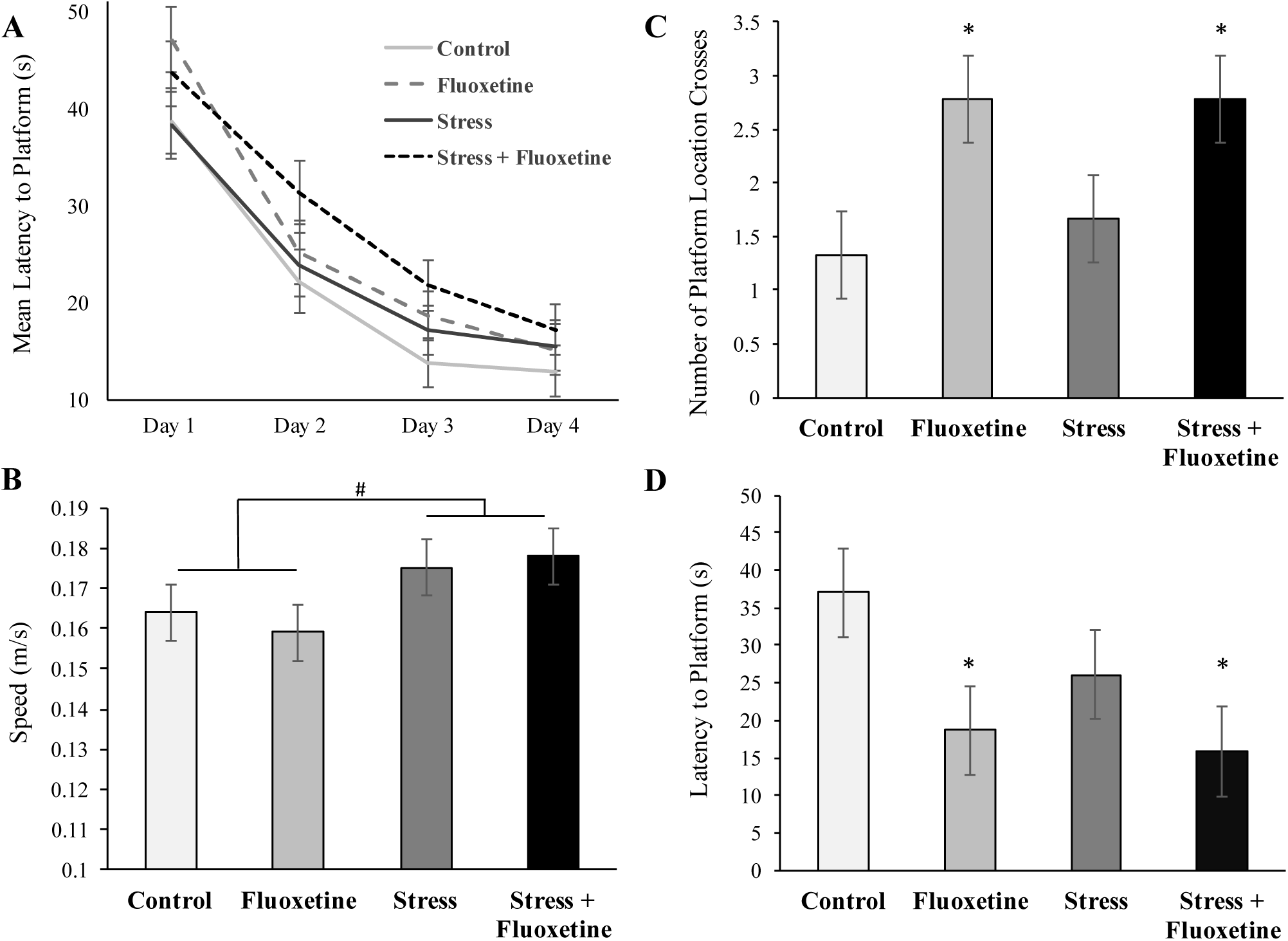
Performance in the Morris water task. Mean latency to reach the platform across 4 days of testing in the MWT (A). Offspring of mice exposed to chronic variable stress traveled with greater speed in the Morris water test (B). Maternal exposure to fluoxetine enhances spatial memory in adult female offspring. In the probe trial, Flx exposure led to an increase in number of crosses of platform’s location (C), and lead to a shorter latency to find platform’s location (D). n=9 per group; # denotes significant main effect of stress; * denotes a significant main effect of fluoxetine; # and * *p* < 0.05. All data are presented as mean ± SEM.

## 4. Discussion

Here, we show that chronic unpredictable maternal stress leads to increased activity and alterations of PPI in the adult female mouse offspring. We also demonstrate that perinatal maternal fluoxetine treatment appears to have little effect on female mice in the behavioural test we have used, potentially having beneficial effects on spatial memory. We further show that the combination of prenatal stress and perinatal Flx exposure did not interact in their effects. We have previously demonstrated no alterations in the maternal behaviour of dams following exposure to stress, Flx, or their combination [32], therefore, behavioural alterations observed in this study are unlikely to be caused by changes in maternal care received by experimental animals.

During early development, the brain is particularly vulnerable to maternal stress; maternal stress is known to lead to short- and long-term changes in offspring brain and behaviour. Maternal stress is linked with higher incidence of neuropsychiatric disorders [76-78], as well as multiple alterations in offspring neuroanatomy and behaviour. The challenge of interpreting the long-term effects of PS is the variation of experimental methods which leads to inconclusive results. Preclinical studies offer no consistent results on the effects of maternal stress on prepulse inhibition and activity levels of adult offspring.

In this study, we found that chronic unpredictable stress administered to C57BL/6 dams from E4 to E18 led to increased activity amongst their female offspring. Mice exposed to PS swim faster in the MWT, and tend to travel longer distances at higher speeds in the EPM. Similar findings have been reported in clinical studies where maternal prenatal stress is linked to childhood attention deficit hyperactivity disorder (ADHD) in their female children [79, 80]. Animal studies also report increased activity in perinatally stressed male [20, 39, 81] and female [25] offspring. In adult animals, chronic stress exposure also leads to increased activity, suggesting a possibility of a common mechanism where early and late-life stress both increase activity levels, however, only male mice have been examined in this regard [82-84].

We found that female mice exposed to PS exhibit increased prepulse inhibition. Lehmann [24] found that prenatally stressed adult female rats similarly show increased PPI. However, the opposite effect of PS on PPI of female rats has also been reported [85]. In humans, PPI alterations are associated with a number of psychiatric disorders [for a review see 86]; increased PPI in females (but not males) is associated with bipolar disorder [87]. Interestingly, PS is linked to bipolar disorder in humans [88]. Like the female mice in our study, human females (but not males) with bipolar disorder have increased PPI [87].

Here, and in previous papers, we show that Flx administration to a dam leads to detectable fluoxetine and norfluoxetine levels in pup brains [28]. Perinatal exposure to Flx has a number of long-term effects on offspring. Studies of male animals show changes in circadian activity, cocaine sensitivity, depressive-like behaviour, anxiety-like behaviour, fear memory, thermal sensitivity, sexual motivation, and copulatory behaviours [for a review see 19]. Few studies examine the effect that exposure to Flx early in development has on behaviour of adult female offspring. At this time, an increase in copulatory behaviour [49], increases and decreases and depressive-like behaviours [43, 70], and a decrease or no change in anxiety-like behaviour have been reported [43, 70, 89].

Independent of stress, perinatal exposure to Flx had only minor effects on the offspring’s behaviour as adults. In the probe trial of MWT, Flx-exposed mice crossed the platform location more times and found the platform location faster than mice not exposed to Flx. Improved MWT performance following perinatal Flx exposure was previously demonstrated in male mice [28], and adolescent male and female rats [90]. Because spatial memory performance is correlated with hippocampal neurogenesis [for a review see 91], the enhancement in the MWT performance may be related to Flx-induced alterations in hippocampal neurogenesis [92-94]. Maternal Flx treatment had no effect on female offspring’s cognitive ability, anxiety, sensorimotor information processing, and exploratory and risk assessment behaviours.

The comparison of our findings to those of males is imperative. The optimal benchmark is our earlier studies in males, as identical stress and Flx treatment protocols were used. Some behavioural changes appear to be sex-dependent. For example, PS has disparate effects on PPI; unlike in males, PPI of female mice is altered by PS [28, 32]. Flx also has sexually-dimorphic effects on anxiety behaviour, having an anxiolytic effect in male, but not female mice [28, 32]. While sex-differences in the effects of stress and Flx are observed, some similarities also exist. PS causes increased activity, while Flx has the potential to improve spatial memory in mice of both sexes [28, 32, 35]. At present, not all behaviours have been studied in mice of both sexes. For example, aggression and circadian behaviours have not yet been examined in female mice exposed to PS and Flx. In males, PS and Flx interact in their effect on these behaviours [32, 35], therefore, examination of aggression and circadian behaviours in females is needed.

While it is becoming clearer that PS, Flx, and their combination lead to different behavioural changes in males and females, the mechanism of such sexually dimorphic changes is not yet understood. A combination of stress-induced changes in maternal glucocorticoids, opioid peptides, and catecholamines may have distinct effects on the development of male vs female offspring brain [95] and may also have the ability to directly alter sexually dimorphic brain areas [44] explaining different outcomes in female and male animals exposed to PS. Furthermore, alterations in the serotonergic system may underlie sex-dependent changes that follow Flx exposure [32, 66, 96]. More research is needed to fully understand the mechanism of sexually dimorphic behavioural changes that follow exposure to PS, Flx, and their combination.

## 5. Conclusion

Due to the high percentage of pregnant women that take Flx, it is important to understand the effect this medication alone, and in combination with maternal stress and depression, has on the offspring. Few studies examine combined effects of stress and Flx, and most of the studies that exist examine only the outcomes of male offspring. In the present study, we report that maternal stress has a number of long-lasting, potentially negative, effects on female mouse offspring. We also report that fluoxetine has very few effects on female mouse offspring, and may even be beneficial due to its potential to improve spatial memory. We further show that, in female offspring, Flx did not worsen nor reverse the effects of maternal stress. Our findings indicate that maternal intake of Flx may have a limited impact on female mouse offspring, however more research is necessary to clearly understand the mechanism of such effects and its translation to humans.

## Acknowledgement of Funding

This work was supported by NSERC Discovery and CIHR Operating grants to RHD, an NSERC and AIHS graduate scholarship to VK and a CIHR summer studentship to SM.

